# Chestnut extract but not sodium salicylate decrease the severity of diarrhea and enterotoxigenic *Escherichia coli* F4 shedding in artificially infected piglets

**DOI:** 10.1101/575662

**Authors:** M. Girard, D. Hu, N. Pradervand, S. Neuenschwander, G. Bee

## Abstract

The development of alternatives to antibiotics is crucial to limiting the incidence of antimicrobial resistance, especially in prophylactic and metaphylactic use to control post-weaning diarrhea (PWD). Feed additives, including bioactive compounds, could be a promising alternative. This study aimed to test two bioactive compounds, sodium salicylate (SA) and a chestnut extract (CE) containing hydrolysable tannins, on the occurrence of PWD. At weaning, 72 piglets were assigned to four treatments that combined two factors: CE supplementation (with 2% of CE (CE+) or without (CE-)) and SA supplementation (with 35 mg/kg BW of SA (SA+) or without (SA-)). Then, 4 days after weaning, all piglets were infected with a suspension at 10^8^ CFU/ml of enterotoxigenic *Escherichia coli* (ETEC F4ac). Each piglet had free access to an electrolyte solution containing, or not, SA. This SA supplementation was administered for 5 days (i.e., from the day of infection (day 0) to 4 days post-infection (day 4). During the 2 weeks post-infection, supplementation with SA had no effect (P > 0.05) on growth performances nor on fecal scores. A significant SA × time interaction (P < 0.01) for fecal scores and the percentage of diarrhea indicated that piglets with SA did not recover faster and did have a second episode of diarrhea. In contrast to SA treatment, inclusion of CE increased (P < 0.05) growth performances and feed intake. In the first week post-infection, CE decreased (P < 0.001) the overall fecal scores, the percentage of piglets with diarrhea, the days in diarrhea, and ETEC shedding in the feces. There was a SA×CE interaction (P < 0.05) for ETEC shedding, suggesting a negative effect of combining SA with CE. This study highlighted that, in contrast to SA, CE could represent a promising alternative to antibiotics immediately after weaning for improving growth performance and reducing PWD.

## Introduction

Post-weaning diarrhea (PWD) is a major enteric disease in pig production that occurs mainly during the first two weeks after weaning. This problem causes substantial economic losses due to high rates of mortality and morbidity, depression of feed intake and growth, and increased costs of medication [1].

Apart from the multifactorial etiology owing to several stressors, including nutritional, social, and environmental changes, PWD is often related to infection with specific pathogens. Enterotoxigenic *Escherichia coli* (ETEC) is a prevalent pathogen involved in PWD [2]. For instance, in Switzerland, 42.5% of weaned pigs with diarrhea were identified as having ETEC [3]. Those bacteria possess fimbriae that adhere to receptors located on the apical side of the enterocytes and secrete enterotoxins, including heat labile toxin (LT) and heat stable toxin (STb), which results in fluid secretion into the intestinal lumen, leading to dehydration and acidosis, which ultimately results in diarrhea [4, 5].

Antibiotics are usually used to treat this type of infection. However, the widespread use of antimicrobials, especially in prophylactic and metaphylactic group treatments of weaned piglets, has led to an increase in microbial resistance. A recent farm survey in Switzerland reported that 70% of enterovirulent *Escherichia coli* (*E. coli*) isolated from pig suffering from diarrhea were resistant to one or more antimicrobials [6]. The field of animal nutrition can contribute to limiting the recrudescence of antimicrobial resistances by exploring nutritional alternatives to antibiotics that prevent microbial infections. Some plants are known as nutraceuticals because they exhibit antimicrobial properties for both susceptible and multiresistant-drug bacteria [7]. It has been demonstrated *in vitro* that polyphenol-rich plants can disrupt ETEC adhesion and inhibit some enterotoxins, including the LT from ETEC [8, 9]. Among polyphenols, hydrolysable tannins from pomegranate and chestnut extract have been shown to impair the growth of bacteria *in vitro*, including *E. coli*, *Listeria monocytogenes*, *Staphylococcus aureus*, *Yersinia enterocolitica, Salmonella enteritidis*, and *Clostridium perfringens* [10, 11].

Pathogen infections provoke inflammatory responses that manifest in loss of appetite and sometimes fever. Previous studies have reported an improvement in performance and a reduction of diarrhea in weaned pigs receiving acetylsalicylic acid, also known as aspirin [12, 13]. Salicylate is an analogue of acetylsalicylic acid and is naturally present in some plants, including sage (*Salvia sp.*) and sweet birch (*Betula lenta*). The primary mode of action of salicylate is to inhibit cyclooxygenase and reduce prostaglandin E2 production, which ultimately reduces the extent of inflammation and stimulates appetite. In addition, salicylate has anti-secretory properties [14].

Thus, the main purpose of the present study was to examine the consequences of supplementation with chestnut extract (CE) containing hydrolysable tannins, either combined or not with sodium salicylate (SA), on growth performance, severity of diarrhea, and bacterial load using a previously established ETEC F4 infection model [15]. An additional goal was to compare two quantitative methods, culturing and quantitative PCR (qPCR), for measuring ETEC shed in the feces. Owing to their antimicrobial and anti-secretory properties, we hypothesize that the inclusion of CE and SA will reduce the severity of diarrhea, with an additive effect when the two compounds are combined.

## Material and methods

### Bacterial strain

The ETEC F4 strain was isolated from a piglet suffering from acute PWD at the Agroscope Posieux (Switzerland) swine facility. The ETEC F4 strain was found to carry the genes for fimbriae F4 (K88ac) and the enterotoxins STb and LT and to grow on an Eosin-Methylene Blue (EMB) agar (CM0069, Oxoid; UK) medium supplemented with 50 µg/ml of rifampicin (rif50). This isolate was stored at −80 °C. The day before infection, the isolate was incubated overnight at 37 °C in Luria-Bertani broth with orbital shaking at 170 rpm. Subsequently, the ETEC F4 inoculum was centrifuged at 6000 rpm for 10 min at room temperature and resuspended in a phosphate buffered saline (PBS) solution to contain approximatively 1 × 10^8^ CFU/ml using the optical density at 600 nm absorbance (Biowave II WPA, LABGENE Scientific SA; Châtel-Saint-Denis, Switzerland).

### Animals and housing

The experiment was conducted in accordance with Swiss guidelines for animal welfare and was approved by the Swiss cantonal veterinary office (approval number: 2016_14_FR). To determine their susceptibility or resistance to ETEC F4ac, genotyping was performed on ear biopsies from week-old piglets [16]. Then, 72 Large White male and female piglets susceptible to ETEC F4ac were selected for the experiment. The piglets were weaned at 26 ± 1 days (mean ± SD) of age, at which time they weighed on average 7.2 ± 1.3 kg (mean ± SD). The animals were housed in pairs in pens of 2.6 m^2^, which were divided into a concrete floor and a galvanized steel floor. Each pen was equipped with a wooden box with straw bedding beneath infrared lamps, a single feeder, a nipple drinker, and a drinking trough. Throughout the experiment, the piglets were kept at an ambient temperature of > 25 °C and had free access to feed, clean water, and a solution of electrolytes (NaCl hypertonic) added to their drinking troughs daily.

### Experimental design, diets and infection challenge

The study was conducted according to a completely randomized 2 × 2 factorial design. The main factors were the CE supplementation (none [CE-] vs. with CE [CE+]), SA supplementation (none [SA-] vs. with SA [SA+]), and the one-way interaction. The resulting four experimental groups were 1) SA-/CE-, piglets receiving a standard diet without supplementation; 2) SA-/CE+, piglets receiving a standard diet with CE and without SA supplementation; 3) SA+/CE-, piglets receiving a standard diet supplemented with SA; and 4) SA+/CE+, piglets receiving a standard diet supplemented with CE and SA.

A total of three farrowing series were necessary to obtain the 72 susceptible piglets (24 piglets each). In each farrowing series, on the day of weaning, piglets were assigned to one of the 4 treatments by balancing the littermates and the weaning body weight (BW). From the day of weaning to the end of the trial, 36 piglets had access to a standard weaning diet (CE-) and 36 to the same diet supplemented with CE (CE+). In the CE+ diet, wheat straw was substituted for with 2% (20 g/kg) of CE: the commercial CE (Silvafeed Nutri P/ENC for Swine, Silvateam; Italy) used contained 45% gallotannins, 9% ellagitannins, and 3.7% gallic acid, for a total of 54% of hydrolysable tannins. The diets were formulated to contain similar nutrient contents, in particular 18% crude protein and 14 MJ/kg of digestible energy, according to the Swiss feed recommendations for pigs [17]. Table 1 shows the compositions and nutrient analyses of the diets. To gradually adapt piglets to the feed, they were given access to a trough containing either CE- or CE+ feed mixed with 100 ml of milk from the day of weaning to one day before infection. Then, 4 days after weaning (i.e., on the day of infection (day 0)), all piglets were orally challenged with an ETEC solution of 5 ml diluted in 50 ml of milk. To ensure that each piglet received the required amount, piglets were kept individually for 10 min. After this time span, piglets which did not ingest the total solution were offered the remaining solution orally via a syringe. Within each feeding group, one-half of the piglets (i.e., 18 fed the CE- diet and 18 fed the CE+ diet) received from days 0‒4 after infection a daily dose of SA (sodium salicylate, ReagentPlus^®^, S3007, Sigma-Aldrich Chemie GmbH; Buchs, Switzerland). The daily dose of SA was dispensed via the electrolyte solution and was based on the total weaning BW of the two piglets in the pen (35 mg/kg BW per day). To maximize the chance of ingestion, the daily doses were offered in a small volume of 500 ml. Based on daily observations, both piglets drank from the solution and in the following mornings, the dispensers were always empty. However, in this setup it was impossible to ensure that each individual piglet ingested the planned dose. The piglets in the SA-groups received a dose of 15 ml of tap water in their electrolyte solution.

**Table 1.**
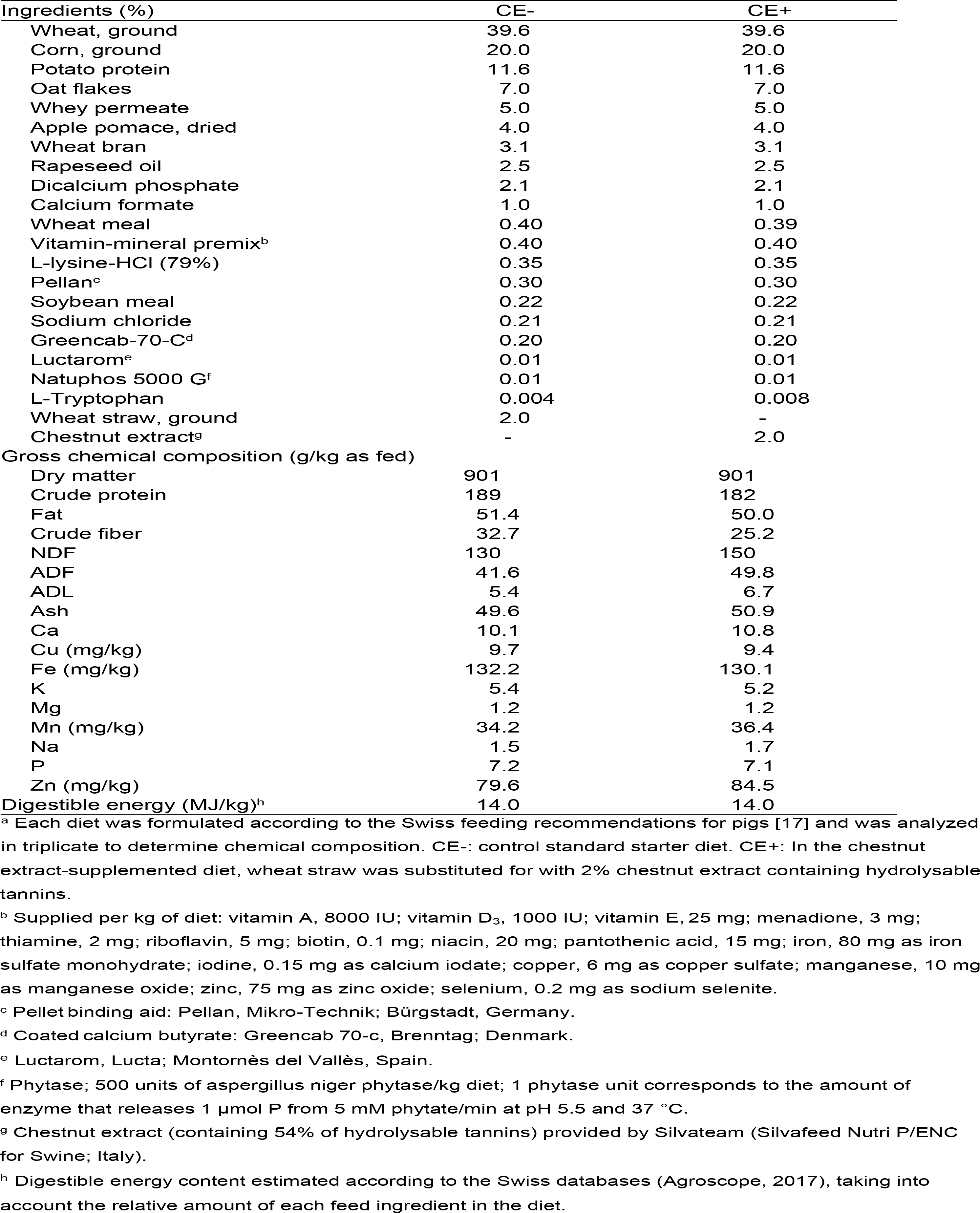
Composition (%) and gross chemical content (g/kg as fed) of the piglets experimental diets^a^

### Measurements, sample collection and analysis

#### Feed analysis

Prior to laboratory analysis, feed samples were ground to pass a 1-mm screen (Brabender mill, Brabender, Duisburg, Germany). Dry matter content was determined by drying feed samples at 105°C for 3h and subsequently analyzed for total ash content by thermogravimetry (prepASH, Precisa Gravimetrics AG, Dietikon, Switzerland). Mineral content of dry ashes was measured according to the European Standard EN 155510:2008 using an inductively coupled plasma optical emission spectrometer (ICP-OES, Optima 7300 DV; Perkin-Elmer, Schwerzenbach, Switzerland). Nitrogen content was analyzed with the Dumas method (ISO 16634-1:2008) and a multiplicative coefficient of 6.25 was applied to calculate crude protein content. Fat content was extracted with petrol ether after an acid hydrolysis (ISO 6492:1999). Crude fiber content was determined gravimetrically (ISO 6865:2000) by incineration of residual ash after acid and alkaline digestions using a fiber analyzer (Fibretherm Gerhard FT-12, C. Gerhardt GmbH & Co. KG, Königswinter, Germany). The NDF, ADF and ADL were analyzed according to standard protocols (ISO 16472:2006 for NDF and ISO 13906:2008 or ADF and ADL) using a fiber analyzer (Fibretherm Gerhard FT-12, C. Gerhardt GmbH & Co. KG, Königswinter, Germany) and were expressed without residual ash and heat-stable amylase and sodium sulfite were used for NDF determination.

### Feed intake per pen, weight and fecal scores

The feed intake per pen was recorded daily during the first week (days 0‒7) and weekly in the second week post-infection (days 8‒14). Individual piglet BW was recorded on day −4 (the day of weaning) and on days 0, 7, and 14. The gain-to-feed ratio per pen was calculated as the sum of the two BW at the end of the period (first week or second week) minus the sum of the 2 BW at the beginning of the respective period within the same pen divided by the total feed intake determined for the pen. The health status of each piglet was closely monitored throughout the experiment. Piglets were observed visually twice a day before infection (days – 4‒ −1), then three times a day during the 5 days after infection (days 0‒4) and then twice a day from days 5‒14. Any unusual behavior, such as apathy, boredom, inappetence, was closely monitored and antimicrobial treatments (according to the veterinarian recommendations) were planned if a piglet showed such behavior. However, in the present study, it did not occur and none of the piglets received antimicrobial treatments. From days 0‒14, every morning at 8h, individual fecal scores were visually evaluated by the same observer for all farrowing series based on the consistency of the feces using the following scores: 1 = dry, molded, or pelleted feces; 2 = creamy, sloppy, or cow-dung appearance; 3 = liquid diarrhea; and 4 = watery diarrhea. Piglets were considered to have diarrhea when the fecal score was ≥ 3. For days 0, 1, 2, 3, 4 and 7, evaluation of fecal score was based on sampled collected from the rectum of each animal. These samples were used to detect the challenge strain using microbial culturing and qPCR techniques. The samples for microbial culturing were directly plated, and the samples for qPCR were frozen at −80 °C for further analysis. For the remaining days, individual fecal scores were evaluated either through direct observation of each piglet defecating or either, when it was not possible, by taking into account the feces on the floor of the pen and whether the anus and the hind legs were irritated, dirty and wet.

### Detecting ETEC F4ac shedding in the feces using microbial culturing and qPCR after DNA extraction

After collection, fecal swab samples were homogenized in 500 μl of sterile PBS for 2 h at room temperature before serial dilution in sterile PBS from 10^0^ to 10^−7^. For each dilution, 4 μl was deposited as a droplet on an EMB rif50 agar plate. The plates were incubated overnight at 37°C, and colonies were counted on each droplet the next morning using binoculars [18]. The mean of the count from two dilutions was used in the results. In addition, those colonies were restricted on a nutrient agar rif50 plate (Becton Dickinson; UK) and cultivated overnight at 37°C to test for the presence of the LT toxin gene using PCR to ensure that the strains growing in the feces were the same as the infection strain from the inoculum.

The feces initially frozen were then freeze dried. DNA was extracted from the dried feces sample using a QIAamp^®^ Fast DNA stool Mini Kit according to the manufacturer’s instructions. A qPCR test was performed. Briefly, the following LT primers (F: 5’-GGCGTTACTATCCTCTCTAT-3’; R: 5’-TGGTCTCGGTCAGATATGT-3’) were used, resulting in a PCR fragment of the 272 base pair [19]. A Bio-Rad CFX96 Touch PCR machine and a KAPA SYBR^®^ FAST qPCR universal kit (KAPA Biosystem) were used. The DNA of the infective ETEC F4ac strain was used as a standard curve. The DNA concentration of standard 1 was 3.1 ng/µl, and serial 1:10 dilutions were conducted for standards 2‒7. In the qPCR, 15 ng of DNA from each feces sample was used as a template. The thermal cycling conditions were 95 °C for 3 min followed by 40 cycles at 95 °C for 10 s, 58 °C for 30 s, and 72 °C for 30s. The melting curve analysis confirmed the primer specificities with the thermal cycling conditions as follows: 95 °C for 10 s and incrementing 0.5 °C per 5 s from 65‒95 °C.

### Statistical analysis

The data for feed intake per pen, BW of the piglets, average daily gain, gain-to-feed ratio per pen and ETEC shedding as determined by plate counting and qPCR were analyzed using linear mixed models in Systat 13 software (Systat Software, Inc.; San Jose, CA, USA). Before analysis, the data for ETEC shedding as determined by plate counting were expressed as log_10_ and data for ETEC shedding as determined by qPCR were transformed as follows: 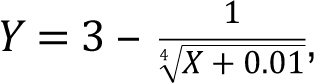 where Y represents the transformed qPCR value and X, the raw qPCR value expressed in ng heat-labile toxin-DNA per g of feces dry matter.

Discrete dependent variables were modeled using R software (R Core Team, 2016): counts for days in diarrhea as quasi-Poisson, ordinal fecal scores as proportional odds logistic regression using generalized estimating equations, and dichotomous responses for percentages of diarrhea as a binary generalized linear mixed model. All models included the effects of SA, CE, farrowing series, time (weeks or days), sex, and the first-order interactions SA × CE, SA × time, and CE × time as fixed effects and either pairs of piglets (for feed intake and gain-to-feed ratio) or piglets, litter and pen as random effects. The BW at day 0 was not included as a covariate for traits related to growth performance because it did not differ between the four treatments at day 0. In general, models were reduced by stepwise exclusion of non-significant interactions and factors (except diet and salicylate) at a P ≤ 0.10. The least squares means of the response variables and Tukey-Kramer pairwise comparisons were computed, and differences were considered as significant if P ≤ 0.05 and as a tendency if P ≤ 0.10.

The correlation between ETEC shedding determined using plate counting and qPCR was estimated by Spearman correlation coefficient. Additionally, the method of Carstensen *et al.*, using the MethComp package of R software, was applied to estimate the variance components used to construct the limits of agreements and the Bland-Altman plots [20]. Before analysis, the data for ETEC shedding as determined by plate counting were expressed as log_10_ and data for ETEC shedding as determined by qPCR as log_10_ (1+X), where X represents the raw qPCR value. The relationship between fecal score and ETEC shedding determined using plate counting or qPCR was analyzed using linear mixed models in Systat 13 software (Systat Software, Inc.; San Jose, CA, USA). As previously, before analysis, the data for ETEC shedding as determined by plate counting were expressed as log_10_ and data for ETEC shedding as determined by qPCR were converted according to the aforementioned fourth root transformation. The models included the effects of fecal score as fixed effect and piglets, litter and pen as random effects.

## Results

### Growth performance and feed intake

Piglets ingested more (P < 0.001) feed and grew faster in the second week post-infection than in the first (Table 2). Piglets from the SA+ groups had performances similar (P > 0.10) to those of piglets from the SA-groups. In contrast, intake per pen in the CE+ groups was on average 20% greater (P = 0.02) than in the CE-groups. Piglets fed the CE+ diet had on average a growth rate 40 g/d greater (P < 0.05) than that of the CE-piglets. Accordingly, the increase in BW in the first and second weeks was more distinct in the CE+ group than in the CE-group (first week: 0.95 vs. 0.60 kg; second week: 1.92 vs. 1.63 kg, respectively [CE × Time interaction: P = 0.001]). Nevertheless, gain-to-feed ratio per pen was not (P > 0.10) affected by the diet.

**Table 2.** Feed intake (g/day per pen), BW (Kg), average daily gain (g/day) and feed efficiency (g/g per pen) from days 0‒7, 8‒14, and 0‒14 post-infection of piglets

| Salicylate<sup>a</sup> | SA- |  | SA+ |  | SEM | P-values<sup>d</sup> |  |  |
| --- | --- | --- | --- | --- | --- | --- | --- | --- |
| Chestnut extract<sup>b</sup> | CE- | CE+ | CE- | CE+ |  | SA | CE | T |
| Body weight (kg)<sup>c</sup> |  |  |  |  |  |  |  |  |
| At day 0 | 7.30 | 7.53 | 7.26 | 7.31 | 0.365 | 0.36 | 0.10 | < 0.001 |
| At day 7 | 8.01 | 8.57 | 7.76 | 8.18 |  |  |  |  |
| At day 14 | 9.67 | 10.47 | 9.37 | 10.14 |  |  |  |  |
| Feed intake per pen (g/day) |  |  |  |  |  |  |  |  |
| Days 0–7 | 415 | 488 | 353 | 458 | 61.7 | 0.22 | 0.02 | < 0.001 |
| Days 8–14 | 827 | 979 | 811 | 887 |  |  |  |  |
| Days 0–14 | 635 | 716 | 546 | 720 | 53.6 | 0.42 | 0.02 | - |
| Average daily gain (g/day) |  |  |  |  |  |  |  |  |
| Days 0–7 | 101 | 149 | 70 | 124 | 21.5 | 0.35 | 0.004 | < 0.001 |
| Days 8–14 | 238 | 273 | 230 | 280 |  |  |  |  |
| Days 0–14 | 169 | 210 | 150 | 202 | 22.5 | 0.52 | 0.04 | - |
| Gain-to-feed per pen (g/g) |  |  |  |  |  |  |  |  |
| Days 0–7 | 0.52 | 0.60 | 0.28 | 0.51 | 0.111 | 0.22 | 0.35 | 0.11 |
| Days 8–14 | 0.62 | 0.59 | 0.59 | 0.59 |  |  |  |  |
| Days 0–14 | 0.55 | 0.59 | 0.55 | 0.56 | 0.039 | 0.62 | 0.59 | - |

<sup>a</sup> SA+: From days 0–4, pigs (N = 36) were offered a daily dose of sodium salicylate (35 mg/kg BW per day; sodium salicylate, ReagentPlus®, S3007, Sigma-Aldrich Chemie GmbH; Buchs, Switzerland). SA-: From days 0–4, pigs (N = 36) were offered a daily dose of tap water.
<sup>b</sup> CE-: Pigs (N = 36) had *ad libitum* access to the control standard starter diet, which was formulated according to the Swiss feeding recommendations for pigs (Agroscope, 2017). CE+: Pigs (N = 36) had *ad libitum* access to the chestnut extract (CE)-supplemented diet, in which wheat straw in the CE- diet was substituted for with 2% CE (Silvafeed Nutri P/ENC for Swine; Italy).
<sup>c</sup> Significant D × T interaction for body weight (P = 0.001).
<sup>d</sup> P-values for the main effect of salicylate supply (SA), chestnut extract supplementation (CE) and time (T).

### Consistency of the feces and percentage of piglets with diarrhea

The consistency of the feces given by the fecal score varied (P < 0.001) over days, with a maximal score on the second or third day post-infection (Fig 1). Supplementing piglets with SA for 5 days post-infection created no (P = 0.70) change in the fecal score, whereas the addition of CE reduced (P < 0.001) the fecal score. In the first week, an increase in fecal score was observed in SA+ piglets during days 5‒7 post-infection, while the fecal scores of SA-piglets decreased progressively over this period (SA × Time interaction: P = 0.01).

**Fig 1.**
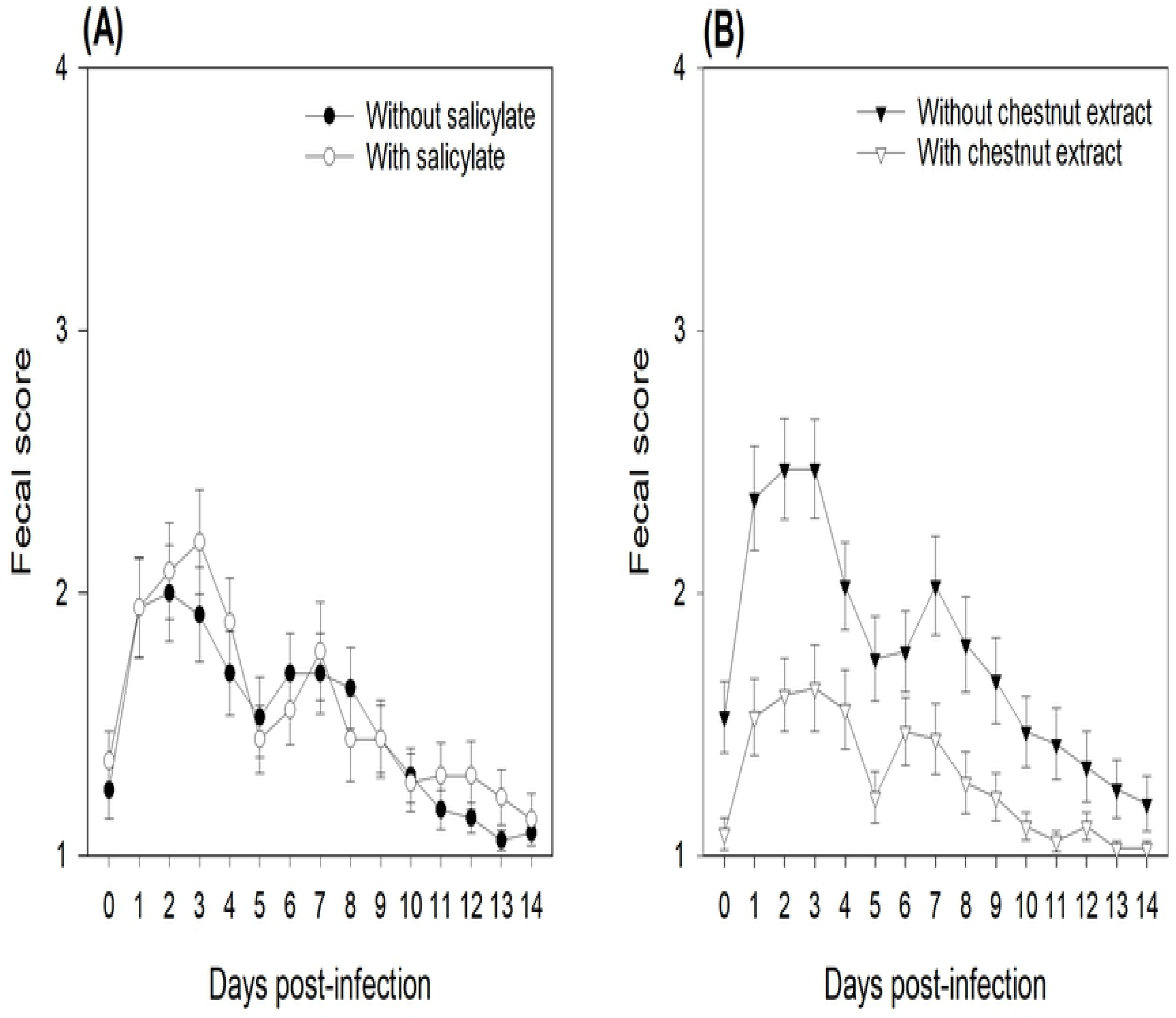
Effects of sodium salicylate supply (A) and chestnut extract supplementation (B) on the fecal scores of piglets assessed from days 0‒14 post-infection. During 14 days post-infection, fecal scores were assessed according to the following score scale: 1 = dry pelleted feces, 2 = cow-dung appearance, 3 = liquid diarrhea, and 4 = watery diarrhea. From days 0‒4, pigs (N = 36) in the SA+ group were offered a daily dose of sodium salicylate (35 mg/kg BW per day, sodium salicylate, ReagentPlus^®^, S3007, Sigma-Aldrich Chemie GmbH; Buchs, Switzerland). From days 0‒4, pigs (N = 36) in the SA-group were offered a daily dose of tap water. Pigs (N = 36) in the CE-group had *ad libitum* access to a control standard starter diet, which was formulated according to the Swiss feeding recommendations for pigs (Agroscope, 2017). Pigs (N = 36) in the CE+ group had *ad libitum* access to the chestnut extract-supplemented diet, in which wheat straw in the CE-diet was substituted for with 2% CE (Silvafeed Nutri P/ENC for Swine; Italy). P-values for the main effects: SA supply: P = 0.70; CE supplementation: P < 0.001; Time: P < 0.001; SA × Time: P = 0.01.

The percentage of piglets with diarrhea was day-dependent (P < 0.001), with the maximum percentage occurring at days 2 and 3 post-infection (Figs 2 A and 2B). The effect of the SA depended on the time (SA × Time interaction: P = 0.003), as in the SA-group, the percentage of piglets with diarrhea increased until day 2, on which 35% of the piglets had diarrhea, and then decreased progressively over days to reach 0% from day 11 onward. In contrast, in the SA+ group, the percentage of piglets with diarrhea increased to 40% at day 2 and then decreased drastically from 35% to 5% between days 4 and 5, when piglets stopped receiving the SA. However, compared to the SA-group, in the SA+ group, from days 5‒7, there was an increase in the percentage of piglets with diarrhea, which decreased again over the following days to stabilize at 4% at the end of the second week post-infection. The addition of CE reduced (P < 0.001) the percentage of piglets with diarrhea. For instance, at 2 days post-infection, 60% of the CE-piglets had diarrhea, but only 16% of the CE+ piglets.

**Fig 2.**
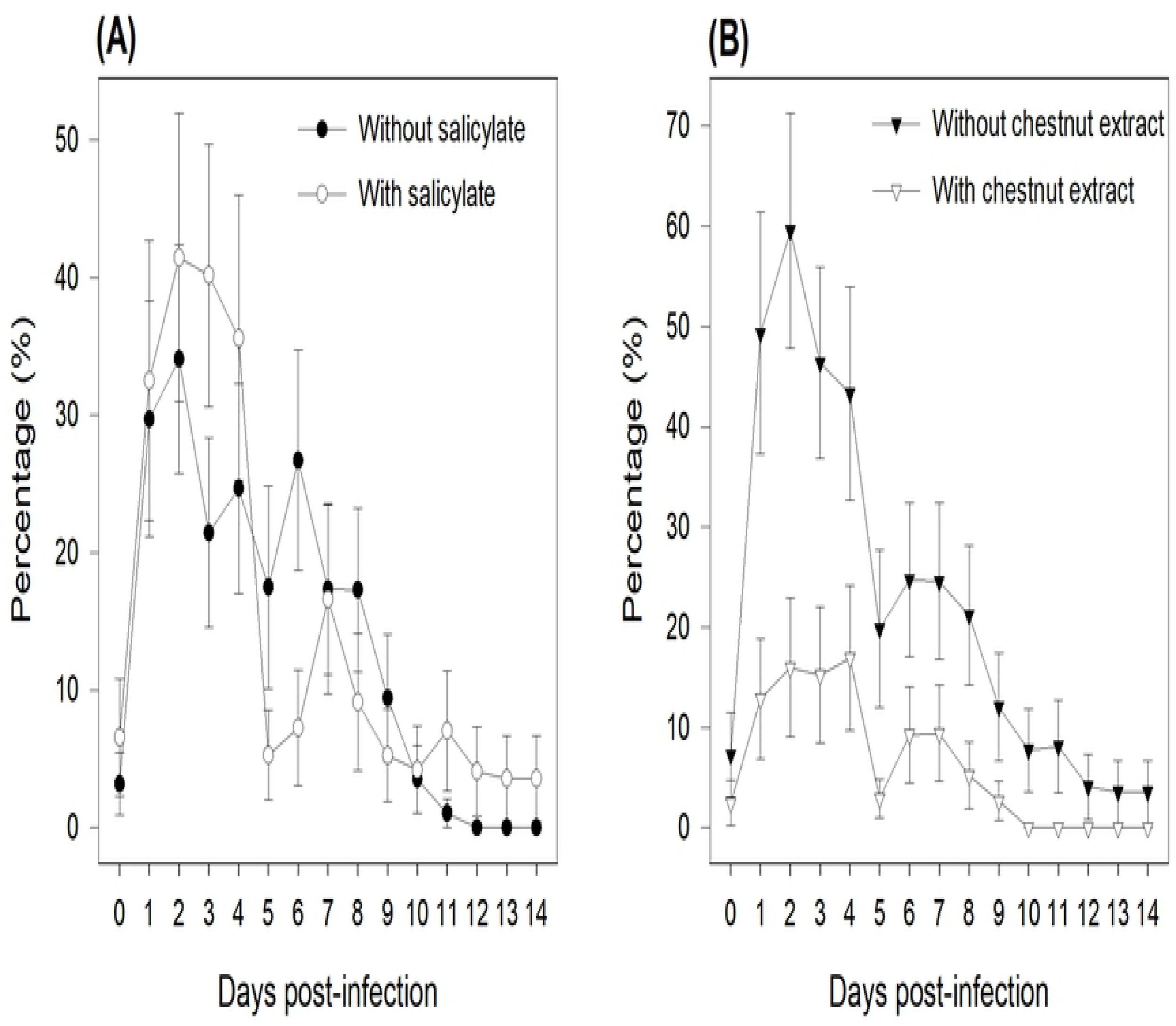
Effects of sodium salicylate supply (A) and chestnut extract supplementation (B) on the percentage of piglets with diarrhea from days 0‒14 post-infection. Fecal scores ≥ 3 were termed diarrhea. From days 0‒4, pigs (N = 36) in the SA+ group were offered a daily dose of sodium salicylate (35 mg/kg BW per day, sodium salicylate, ReagentPlus^®^, S3007, Sigma-Aldrich Chemie GmbH; Buchs, Switzerland). From days 0‒4, pigs (N = 36) in the SA-group were offered a daily dose of tap water. Pigs (N = 36) in the CE-group had *ad libitum* access to a control standard starter diet, which was formulated according to the Swiss feeding recommendations for pigs (Agroscope, 2017). Pigs (N = 36) in the CE+ group had *ad libitum* access to the chestnut extract (CE)-supplemented diet, in which wheat straw in the CE-diet was substituted for with 2% CE (Silvafeed Nutri P/ENC for Swine; Italy). P-values for the main effects: SA supply: P = 0.72; CE supplementation: P < 0.001; Time: P < 0.001; SA × Time: P = 0.003.

## Days in diarrhea

The SA supplementation had no effect on the number of days in diarrhea in the first and second weeks post-infection (Table 3). In the first week post-infection, only the SA-/CE+ piglets tended to have fewer days in diarrhea than the SA-/CE-ones, whereas no difference was observed between the SA+/CE+ and the SA+/CE-groups (SA × CE interaction; P = 0.06). In the second week post-infection, piglets fed the CE+ diet had fewer (P < 0.001) days in diarrhea than those on the CE-diet. Finally, over the two experimental weeks post-infection, CE+ piglets had, on average, diarrhea (i.e., fecal scores ≥ 3) for 1.4 days (P < 0.001), whereas CE-piglets had 4.2 days in diarrhea.

**Table 3.** Number of days in diarrhea (i.e., fecal score ≥ 3) in the two weeks after infection in piglets

| Salicylate<sup>a</sup> | SA- |  | SA+ |  | SEM | P-values<sup>c</sup> |  |
| --- | --- | --- | --- | --- | --- | --- | --- |
| Chestnut extract<sup>b</sup> | CE- | CE+ | CE- | CE+ |  | SA | CE |
| Days 0–7<sup>d</sup> | 3.3<sup>y</sup> | 0.7<sup>x</sup> | 3.0<sup>y</sup> | 1.8<sup>xy</sup> | 0.61 | 0.41 | < 0.001 |
| Days 8–14 | 0.9 | 0.1 | 1.3 | 0.2 | 0.35 | 0.44 | < 0.001 |
| Days 0–14 | 4.1 | 0.8 | 4.3 | 2.0 | 0.88 | 0.35 | < 0.001 |

<sup>x,y</sup> Rows carrying no common superscript tended to differ at $P < 0.10$ .
<sup>a</sup> SA+: From days 0–4, pigs (N = 36) were offered a daily dose of sodium salicylate (35 mg/kg BW per day; sodium salicylate, ReagentPlus®, S3007, Sigma-Aldrich Chemie GmbH; Buchs, Switzerland). SA-: From days 0–4, pigs (N = 36) were offered a daily dose of tap water.
<sup>b</sup> CE-: Pigs (N = 36) had *ad libitum* access to the control standard starter diet, which was formulated according to the Swiss feeding recommendations for pigs (Agroscope, 2017). CE+: Pigs (N = 36) had *ad libitum* access to the chestnut extract (CE)-supplemented diet, in which wheat straw in the CE- diet was substituted for with 2% CE (Silvafeed Nutri P/ENC for Swine; Italy).
<sup>c</sup> P-values for the main effect of salicylate supply (SA), chestnut extract supplementation (CE)
<sup>d</sup> Tendency for SA × CE interaction ( $P = 0.06$ ).

### Fecal ETEC shedding as determined using microbial culturing and using qPCR

ETEC shedding in the feces was determined using both culturing and qPCR. Using either method, the extent of ETEC shedding was not affected by the SA supply, but it was lower (P = 0.01) in piglets fed the CE+ diet than in those fed the CE-diet (Figs 3 and 4). There was a time effect (P ≤ 0.05), as ETEC shedding increased at day 1 post-infection. The results obtained using qPCR also revealed a CE × time interaction (P = 0.02) due to constant shedding around a value of 1.5 over time for the CE+ piglets (except at day 1), while the excretion profile increased to a maximum of 1.94 at day 3 post-infection and then decreased from days 3‒7 for CE-piglets (Fig 4).

**Fig 3.**
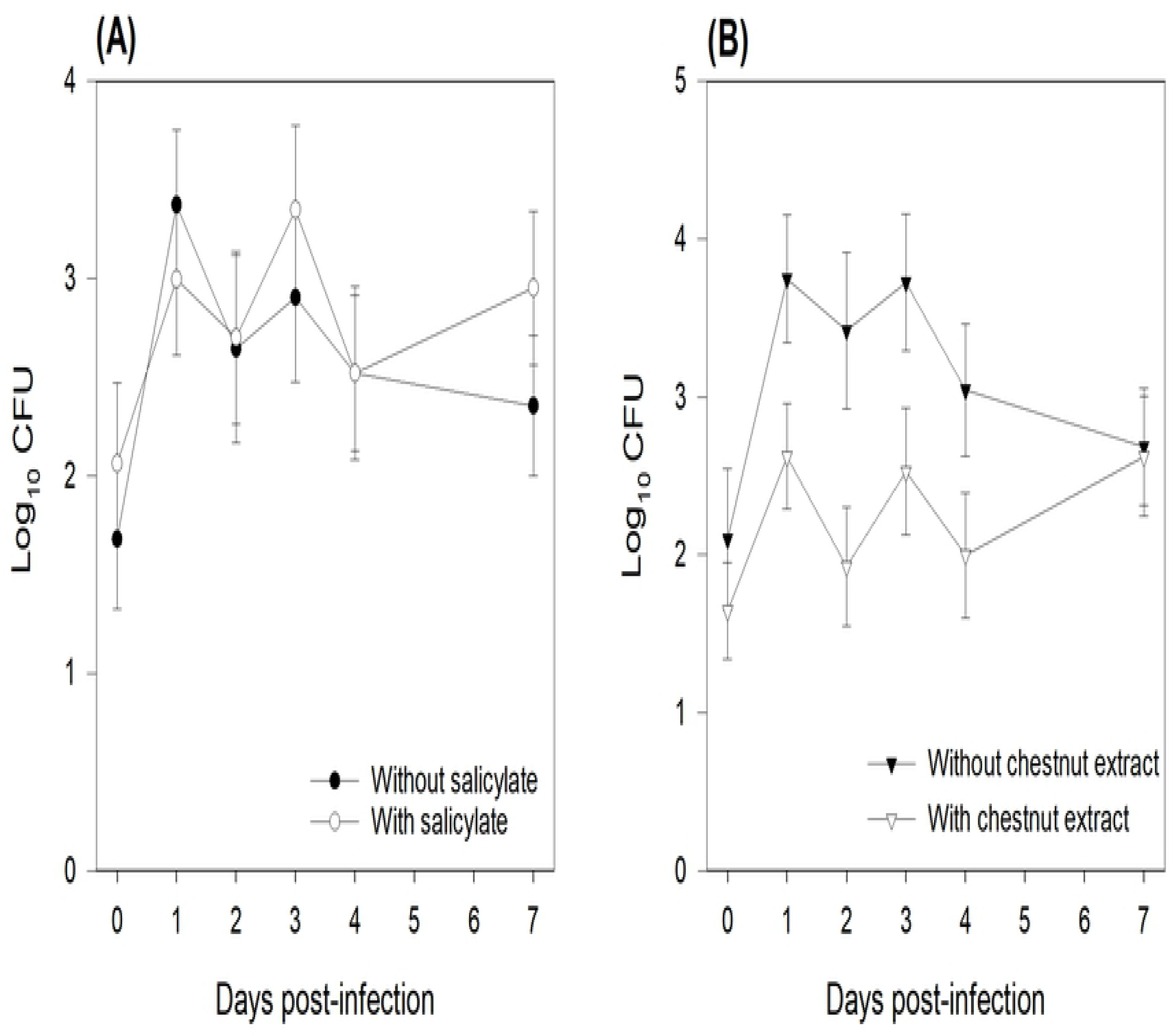
Effects of sodium salicylate supply (A) and chestnut extract supplementation (B) on *Enterotoxigenic Escherichia coli* F4 shedding as determined using plate counting in feces samples collected from all piglets at days 0, 1, 2, 3, 4 and 7 post-infection. Data are expressed as log_10_ (Colony Formant Units). From days 0‒4, pigs (N = 36) in the SA+ group were offered a daily dose of sodium salicylate (35 mg/kg BW per day, sodium salicylate, ReagentPlus^®^, S3007, Sigma-Aldrich Chemie GmbH; Buchs, Switzerland). From days 0‒4, pigs (N = 36) in the SA-group were offered a daily dose of tap water. Pigs (N = 36) in the CE-group had *ad libitum* access to a control standard starter diet, which was formulated according to the Swiss feeding recommendations for pigs (Agroscope, 2017). Pigs (N = 36) in the CE+ group had *ad libitum* access to the chestnut extract (CE)-supplemented diet, in which wheat straw in the CE-diet was substituted for with 2% CE (Silvafeed Nutri P/ENC for Swine; Italy). P-values for the main effects: SA supply: P = 0.51; CE supplementation: P = 0.01; Time: P = 0.02.

**Fig 4.**
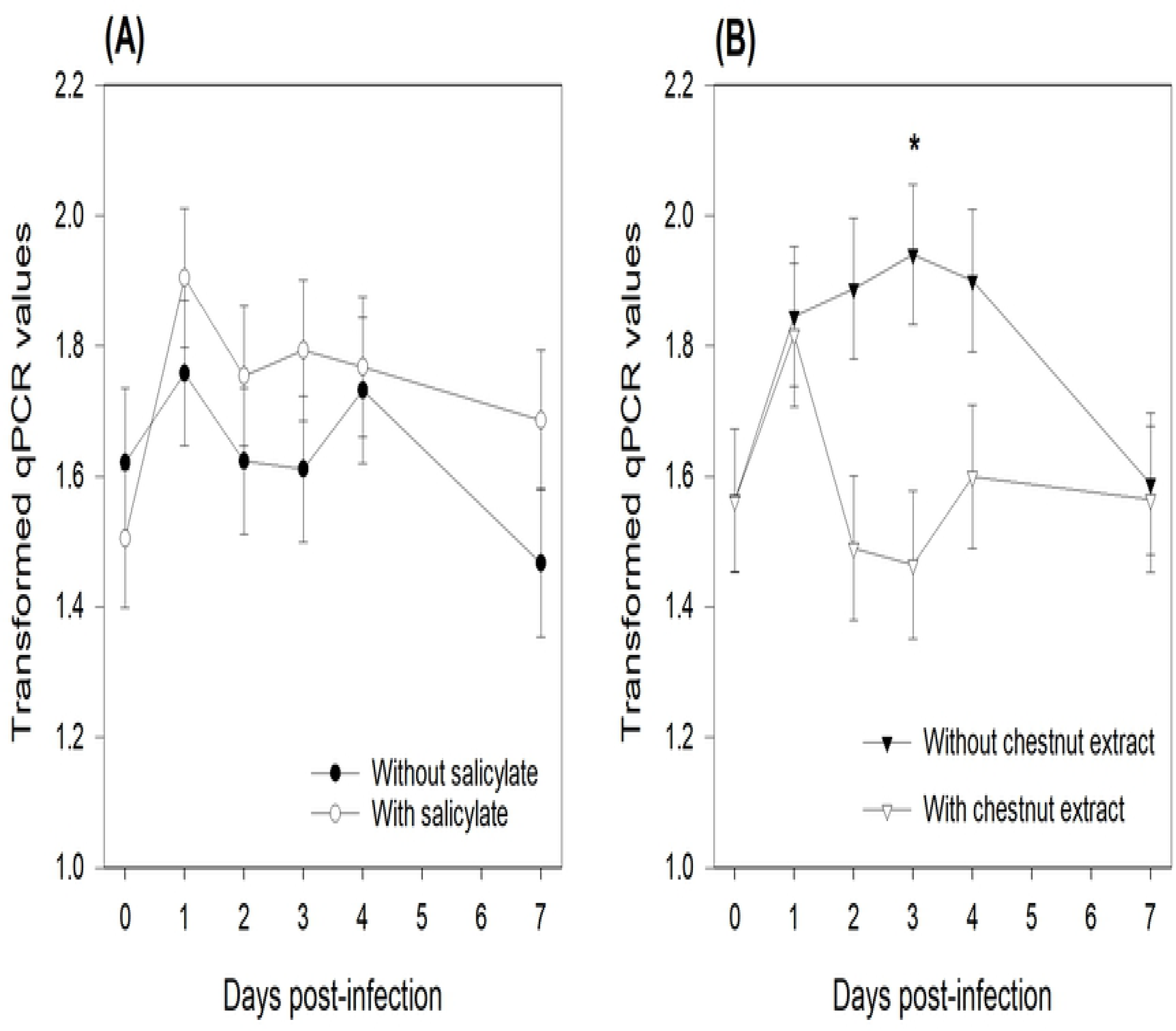
Effect of sodium salicylate supply (A) and chestnut extract supplementation (B) on heat-labile toxin (LT) gene abundance as determined using qPCR in feces samples collected from all piglets at days 0, 1, 2, 3, 4, and 7 post-infection. Data are presented as transformed qPCR values according to: 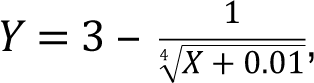 where X represents the ng heat-labile <toxin-DNA per g of feces dry matter. From days 0‒4, pigs (N = 36) in the SA+ group were offered a daily dose of sodium salicylate (35 mg/kg BW per day, sodium salicylate, ReagentPlus^®^, S3007, Sigma-Aldrich Chemie GmbH; Buchs, Switzerland). From days 0‒4, pigs (N = 36) in the SA-group were offered a daily dose of tap water. Pigs (N = 36) in the CE-group had *ad libitum* access to a control standard starter diet, which was formulated according to the Swiss feeding recommendations for pigs (Agroscope, 2017). Pigs (N = 36) in the CE+ group had *ad libitum* access to the chestnut extract (CE)-supplemented diet, in which wheat straw in the CE-diet was substituted for with 2% CE (Silvafeed Nutri P/ENC for Swine; Italy). P-values for the main effects: SA supply: P = 0.25; CE supplementation: P = 0.01; Time: P = 0.001; CE × Time: P = 0.02. * marks differences between CE- and CE+ at P < 0.05 within the same day.

The results obtained using the plate-count technique showed that independent of days post-infection, ETEC shedding was lower in the SA-/CE+ piglets than in the SA-/CE-ones, with intermediate values for the SA+/CE+ and the SA+/CE-piglets (SA × CE interaction: P = 0.03; Fig 5, black bars). Using the qPCR technique showed that SA-/CE+ piglets excreted numerically less ETEC in their feces than did piglets in the three other groups, however it was no significant (SA × CE interaction: P = 0.19; Fig 5, white bars).

**Fig 5.**
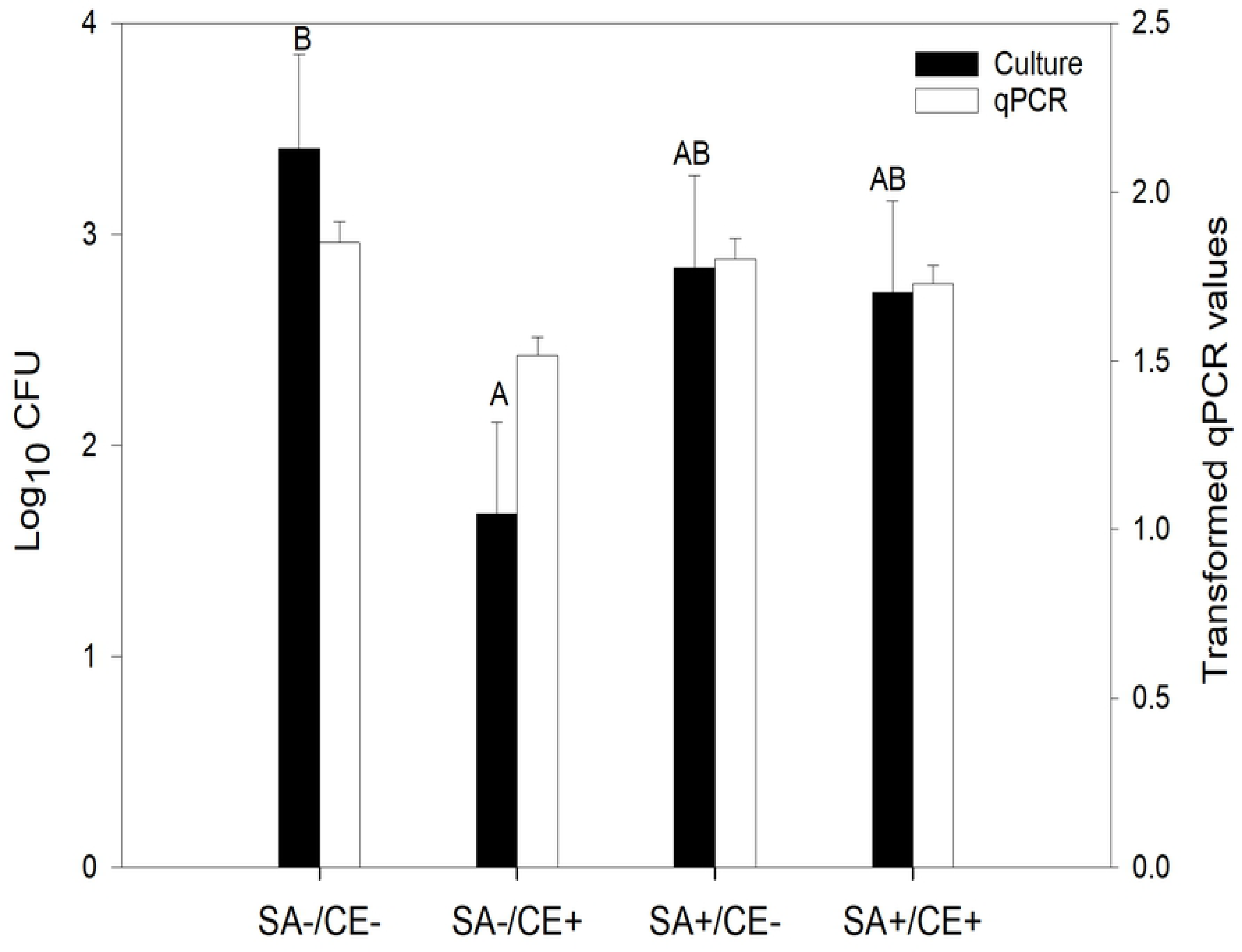
*Enterotoxigenic Escherichia coli* F4 shedding determined using culturing (black bars) and qPCR (white bars) in feces samples from piglets receiving a standard diet without chestnut extract (CE) nor sodium salicylate (SA) supplementation (SA-/CE-; N = 18), from piglets receiving a standard diet with CE and without SA supplementation (SA-/CE+; N = 18), from piglets receiving a standard diet supplemented with SA (SA+/CE-; N = 18), and from piglets receiving a standard diet supplemented with both CE and SA (SA+/CE+; N = 18). From days 0‒4 post-infection, pigs from the SA+/CE-and SA+/CE+ groups were offered a daily dose of sodium salicylate (35 mg/kg BW per day, sodium salicylate, ReagentPlus^®^, S3007, Sigma-Aldrich Chemie GmbH; Buchs, Switzerland). From days 0‒4 post-infection, pigs from the SA-/CE-and SA-/CE+ groups were offered a daily dose of tap water. The CE-diet was a control standard starter diet, which was formulated according to the Swiss feeding recommendations for pigs (Agroscope, 2017). The CE+ diet was a CE-supplemented diet in which the wheat straw in the CE-diet was substituted for with 2% CE (Silvafeed Nutri P/ENC for Swine; Italy). Data are expressed as log_10_ (Colony Formant Units) for culturing and as transformed qPCR values according to: 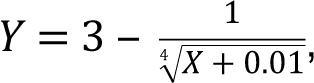 where X represents the ng heat-labile toxin-DNA per g of feces dry matter for qPCR. P-values for SA × CE: P = 0.03 (culturing) and P= 0.19 for qPCR). ^A,B^ Bars carrying no common superscript differ at P < 0.05 for the culturing method.

The ETEC shedding results obtained using the plate culturing and qPCR methods correlated weakly (Spearman rank coefficient = 0.53; P < 0.001; S1 Fig). In addition, the Bland-Altman analysis of the range standardized log data according to Carstensen *et al.* [20] showed limits of agreement far beyond acceptable values (data not shown). Regardless the treatments, the fecal score positively correlated with ETEC shedding with coefficients of 0.62 and 0.11 for plate culturing and qPCR methods respectively (P < 0.001; S2 Fig). Throughout the duration of the experiment, piglets in the SA-/CE+ group never developed watery diarrhea, which would be fecal scores of 4.

## Discussion

The present experiment aimed to elucidate whether the addition of 2% CE with or without SA reduced the severity of diarrhea in piglets artificially infected with ETEC F4ac. A previous experiment successfully established the induction of diarrhea using the same ETEC F4ac infection model [15]. In the present study, a secondary goal was to compare the extent of ETEC shedding as determined using either culturing or qPCR. The results showed a weak positive correlation between the two methods. Lindsay *et al.* reported a slightly greater Spearman’s coefficient (r = 0.61) between the two methods in human volunteers infected with ETEC [21]. The qPCR method amplified the DNA of both dead and living ETEC, while only the living ETEC grew on the culture plates, which could partially explain the discrepancy between the two methods.

### Effects of sodium salicylate

In plants, salicylate belongs to a diverse group of plant phenolics and is a major plant hormone [22]. Some authors have shown a greater average daily gain in piglets either uninfected or infected with ETEC F4 and receiving aspirin, an analogue of salicylate [12, 13]. Van der Klis showed that, owing to its anti-inflammatory properties, SA supplementation via drinking water eliminated the negative effects of *Clostridium* infection on the BW gain of broilers [23]. However, in the present experiment, the same dose of SA offered via electrolyte solution did not improve the growth performance of artificially infected piglets. The SA has also been reported to exert antibacterial effects by affecting bacterial cell growth [24]. However, in the present study, the use of SA did not reduce ETEC F4 shedding in the feces. In addition, SA supplementation reduced neither the overall fecal score nor the percentage of piglets with diarrhea. Furthermore, the significant effect of SA on fecal score and percentage of piglets with diarrhea over time indicates that piglets receiving SA for 5 days post-infection did not recover faster after an episode of diarrhea and had a second episode of diarrhea in the second week post-infection. This result contradicts those showing a supposedly anti-secretory property of SA. Wise *et al.* observed that an infusion of SA decreased fluid accumulation in intestinal loops incubated with *E. coli* ST enterotoxin in calves [14].

### Effects of chestnut extract supplementation

Unlike with SA, piglets supplemented with 2% CE coped better with ETEC infection than piglets who received a standard weaning diet. First, piglets developed less diarrhea with CE supplementation, which is reflected in their decreased fecal scores, percentage of piglets with diarrhea and days in diarrhea during the two weeks post-infection in comparison to piglets fed the CE-diet. The results of the present study were more promising than those of a previous study that supplemented the same hydrolysable tannins CE at a dose of 1% [15]. The development of coliform diarrhea depends on the ability of ETEC to adhere to and colonize the intestinal epithelium. An indirect way to check for the presence of ETEC is to measure ETEC shedding in the feces. Thus, an increase in ETEC shedding in the feces might be the consequence of greater bacterial colonization due to the peristaltic clearance of ETEC [25]. In the present experiment, a positive correlation was observed between diarrhea occurrence as assessed using fecal score and ETEC shedding determined using plate counting or qPCR, which means that the greater the fecal score, the greater the shedding. However a high variability among piglets in the same group exists. Some piglets presented no clinical signs of diarrhea but excreted a high amount of ETEC, which might translate into an improved immunity against the infection. In contrast, other piglets had diarrhea without shedding ETEC, which might have been due to the development of nutritional or viral induced types of diarrhea. Nevertheless, none of the 18 SA-/CE+ piglets developed watery diarrhea (fecal score = 4), and in general, piglets supplemented with CE diet shed less ETEC in their feces. This result agrees with those of previous experiments in which infected or non-infected piglets fed various types of tannin extract excreted less ETEC or F18 verotoxigenic *E. coli* [26-28]. This decreased bacterial shedding might be the result of a bactericidal and/or bacteriostatic effect of tannins on ETEC F4ac, as previously observed *in vitro* [29, 30], and/or a decrease in bacterial adhesion to the intestinal epithelium [27]. A previous study on uropathogenic *E. coli* reported that cranberry tannins were able to decrease *in vitro* the adhesion forces between the bacteria and a probe surface and to alter the conformation of the surface macromolecules on *E. coli* [31]. Nevertheless, besides a positive effect of tannins on reducing diarrhea, previous experiments have failed to show a positive effect of tannins on ETEC shedding [15, 28]. The lack of effects of tannins observed in previous studies is probably related to differences in the bioactivity of tannins according to the dose (1% CE, as previously administered, was not sufficiently high compared to 2% in the present study), as well as to the chemical structure of tannins [32]. However, caution should be taken in using hydrolysable tannins in experiments because up to now, it has been unclear whether the effects of tannins are due to the hydrolysable tannins and/or to their metabolites (ellagic acid and urolithins), as hydrolysable tannins can be hydrolyzed both in the stomach and the gut [33].

In the present study, supplementation with 2% CE for 19 days beginning at weaning improved feed intake and average daily BW gain. Even though tannins are often considered to be an anti-nutritional factor, the present results confirmed those of previous studies in which tannins did not negatively affect animal performance [34, 35]. The improved performance of the CE+ piglets is probably the consequence of a reduction in the occurrence of diarrhea in piglets fed CE, because the CE did not improve feed efficiency.

In the present study, the substitution of 2% wheat straw with CE minimized the differences in terms of ingredients and chemical compositions between the two diets. However, a slight difference in terms of fiber contents can be noticed between the two diets. The diet including CE contained slightly less crude fiber and more NDF, ADF, and ADL than the standard diet. The source and chemical characteristics of fiber components affect the digesta retention time and the fermentation profile in the gut microbiota. Increasing the content of less-fermentable fiber is associated with a reduction in *E. coli* count in the feces [36]. In a recent study, Nepomuceno *et al.* observed a linear decrease in the occurrence of diarrhea when NDF content was increased from 8.5 to16.5% [37]. Hence, in addition to the hydrolysable tannins, the 2% unit greater NDF content in the diet containing CE could have partially helped in reducing both ETEC shedding and the occurrence of diarrhea.

### Effects of the combination sodium salicylate and chestnut extract

The present experiment showed a slight effect of combining SA and CE on the number of days in diarrhea (tendency) in the first week post-infection and on ETEC shedding determined using plate counting. Interestingly, it seemed that SA erased the positive effect of CE, as CE supplementation reduced (or tended to reduce) ETEC shedding and days in diarrhea only when they were fed alone and not combined with SA. An increase in the level of intrinsic antibiotic resistance has sometimes been observed when bacteria grow in the presence of salicylate [38]. In *E. coli*, salicylate has increased the transcription of the multiple antibiotic resistance operon *marRAB* [39]. In addition, the activation of the operon has decreased the accumulation of antibiotics by reducing the synthesis of porins and has increased the production of multidrug efflux pump [39, 40]. Such a mechanism could also be considered if hydrolysable tannins or their metabolites entered the bacteria. However, further investigations are needed to elucidate the effects of salicylate on tannins.

## Conclusion

The present study demonstrated that unlike SA, 2% hydrolysable tannins CE could be beneficial directly at weaning to reduce PWD and thereby enhance piglet growth. Nevertheless, the effect of CE should to be validated in larger-scale trials with more complex conditions. Owing to the ability of hydrolysable tannins to affect various pathogens, it would also be interesting to investigate whether this CE extract is able to affect other pathogens *in vivo*. Finally, the effects of SA on tannins deserve further investigation.

## Acknowledgements

The authors gratefully acknowledge and thank the skilled technical assistance at the experimental piggery (Mr. Guy Maikoff and his team) and the analytic chemistry (Mr. Sébastien Dubois and his team) and microbiology (Dr. Nicolas Pradervand and his team) departments of Agroscope in Posieux. We also thank Dr. Werner Luginbühl of ChemStat (Chemometrik und Statistik, Berne) for his assistance with statistical analysis and Silvateam (Italy) for providing the chestnut extract.

## supporting information

**S1 Fig. Relationship between Enterotoxigenic *Escherichia coli* F4 shedding in feces determined by plate counting and by qPCR (P< 0.001; Spearman rank coefficient = 0.53) in piglets receiving a standard diet without chestnut extract (CE) nor sodium salicylate (SA) supplementation (SA-/CE-), piglets receiving a standard diet with CE and without SA supplementation (SA-/CE+), piglets receiving a standard diet supplemented with SA (SA+/CE-), and piglets receiving a standard diet supplemented with both CE and SA (SA+/CE+).**Pigs of the SA+/CE- and SA+/CE+ groups were offered from days 0 to 4 post-infection a daily dose of sodium salicylate (35 mg/kg BW per day, sodium salicylate, ReagentPlus^®^, S3007, Sigma-Aldrich Chemie GmbH, Buchs, Switzerland). Pigs of the SA-/CE- and SA-/CE+ groups were offered from days 0 to 4 a daily dose of tap water. The CE-diet was a control standard starter diet, which was formulated according to the Swiss feeding recommendations for pigs (Agroscope, 2017). The CE+ diet was a CE-supplemented diet where wheat straw in the CE-diet was substituted with 2% CE (Silvafeed Nutri P/ENC for Swine, Italy).Data are expressed as Log_10_ (Colony Formant Units) for microbial culture and as Log_10_ (1+X) where X represents the raw qPCR data expressed in ng heat-labile toxin-DNA per g of feces dry matter for qPCR.

**S2 Fig. Relationship between Enterotoxigenic *Escherichia coli* F4 shedding in feces determined by plate counting (A) or by qPCR (B) and fecal score (1 and 2: no diarrhea; 3 and 4: diarrhea) in receiving a standard diet without chestnut extract (CE) nor sodium salicylate (SA) supplementation (SA-/CE-), piglets receiving a standard diet with CE and without SA supplementation (SA-/CE+), piglets receiving a standard diet supplemented with SA (SA+/CE-), and piglets receiving a standard diet supplemented with both CE and SA (SA+/CE+).**Pigs of the SA+/CE- and SA+/CE+ groups were offered from days 0 to 4 post-infection a daily dose of sodium salicylate (35 mg/kg BW per day, sodium salicylate, ReagentPlus^®^, S3007, Sigma-Aldrich Chemie GmbH, Buchs, Switzerland). Pigs of the SA-/CE- and SA-/CE+ groups were offered from days 0 to 4 a daily dose of tap water. The CE-diet was a control standard starter diet, which was formulated according to the Swiss feeding recommendations for pigs (Agroscope, 2017). The CE+ diet was a CE-supplemented diet where wheat straw in the CE-diet was substituted with 2% CE (Silvafeed Nutri P/ENC for Swine, Italy). Data are expressed as Log_10_ (Colony Formant Units) for microbial culture and as 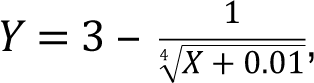 where Y represents the transformed qPCR value and X, the raw qPCR value expressed in ng heat-labile toxin-DNA per g of feces dry matter for qPCR. Black bold lines represent the mean values and black dot indicate samples below or above the 10th and 90th percentiles respectively. (P< 0.001 for plate counting and for qPCR).

